# Integration of genome-wide association studies and gene coexpression networks unveils promising soybean resistance genes against five common fungal pathogens

**DOI:** 10.1101/2021.08.31.458388

**Authors:** Fabricio Almeida-Silva, Thiago M. Venancio

## Abstract

Soybean is one of the most important legume crops worldwide. However, soybean yield is dramatically affected by fungal diseases, leading to economic losses of billions of dollars yearly. Here, we integrated publicly available genome-wide association studies and transcriptomic data to prioritize candidate genes associated with resistance to *Cadophora gregata, Fusarium graminearum, Fusarium virguliforme, Macrophomina phaseolina*, and *Phakopsora pachyrhizi*. We identified 188, 56, 11, 8, and 3 high-confidence candidates for resistance to *F. virguliforme, F. graminearum, C. gregata, M. phaseolina and P. pachyrhizi*, respectively. The prioritized candidate genes are highly conserved in the pangenome of cultivated soybeans and are heavily biased towards fungal species-specific defense response. The vast majority of the prioritized candidate resistance genes are related to plant immunity processes, such as recognition, signaling, oxidative stress, systemic acquired resistance, and physical defense. Based on the number of resistance alleles, we selected the five most resistant accessions against each fungal species in the soybean USDA germplasm. Interestingly, the most resistant accessions do not reach the maximum theoretical resistance potential. Hence, they can be further improved to increase resistance in breeding programs or through genetic engineering. Finally, the coexpression network generated here is available in a user-friendly web application (https://soyfungigcn.venanciogroup.uenf.br/) and an R/Shiny package (https://github.com/almeidasilvaf/SoyFungiGCN) that serve as a public resource to explore soybean-pathogenic fungi interactions at the transcriptional level.

## 1 Introduction

Soybean (*Glycine max* (L.) Merr.) is a major legume crop worldwide, contributing to global food security and economy. However, soybean yield is significantly affected by diseases, with an estimated economic loss of 95.8 billion dollars from 1996 to 2006 in the US (Bandara *et al*., 2020). Most of the yield loss has been linked to foliar and stem/root diseases, which are mostly caused by phytopathogenic fungi (Bandara *et al*., 2020). Fungal diseases, such as sudden death syndrome, Fusarium wilt, brown stem rot and asian rust, can impact soybean crops through leaf damage, necrosis, chlorosis, and death (Pandey *et al*., 2011; Rincker *et al*., 2016; Bandara *et al*., 2020).

Over the past decade, several genome-wide association studies (GWAS) have uncovered multiple single-nucleotide polymorphisms (SNPs) associated with resistance to pathogenic fungi in soybean populations (Iquira *et al*., 2015; Zhang *et al*., 2015, 2019; Chang *et al*., 2016; Rincker *et al*., 2016; Kandel *et al*., 2018; Sun *et al*., 2020). Nevertheless, GWAS often fail to accurately pinpoint the causative genes (Baxter, 2020). GWAS limitations are particularly challenging for self-pollinating plants (*e.g*., soybean) because of limited recombination and strong linkage disequilibrium between causative and non-causative variants (Michno *et al*., 2020). Such limitations ultimately lead to large genetic intervals with several genes, hindering causative gene identification. Because of the exponential accumulation of genomic and transcriptomic data in public databases (Deshmukh *et al*., 2014; Schaefer *et al*., 2018; Wen *et al*., 2018; Baker *et al*., 2019; Schwartz, 2020), integrative analyses to prioritize candidate genes have become a promising approach. This strategy consists in investigating the transcriptional patterns of all the genes near a significant SNP. Hence, the combination of multiple sources of evidence can result in richer and narrower sets of high-confidence candidate genes for downstream experimental validation towards biotechnological applications.

Here, we integrated multiple publicly available RNA-seq and GWAS datasets to identify high-confidence candidate genes for resistance to five phytopathogenic fungi. The prioritized resistance genes are species-specific and highly conserved in the pangenome of cultivated soybeans. The candidate resistance genes against each species are involved in various immunity-related processes, such as recognition, signaling, oxidative stress, and apoptosis. Finally, we highlighted the five most resistant accessions against each fungal species in the USDA germplasm, uncovering important information for breeding programs and genetic engineering initiatives. Finally, the coexpression network resulting from this work was also made available as a publicly available web application (https://soyfungigcn.venanciogroup.uenf.br/) and R/Shiny package (https://github.com/almeidasilvaf/SoyFungiGCN).

## 2 Materials and Methods

### 2.1 Curation of resistance-associated SNPs

SNPs that contribute to resistance against phytopathogenic fungi were manually curated from the scientific literature (Table 1; Supplementary Table S1). SNPs that were identified using the Gmax_a1.v1 genome were converted to their corresponding sites in the Gmax_a2.v1 assembly using the .vcf files for both assemblies available at Soybase (Brown *et al*., 2020).

**Table 1.** GWAS included in this work.

| Reference | Pathogen | Resistance SNPs |
| --- | --- | --- |
| (Zhang *et al.*, 2019) | *F. graminearum* | 12 |
| (Bao *et al.*, 2015) | *F. virguliforme* | 8 |
| (Chang *et al.*, 2016) | *C. gregata* / *F. virguliforme* / *P. pachyrhizi* | 2 / 1 / 2 |
| (Zhang *et al.*, 2015) | *F. virguliforme* | 32 |
| (Swaminathan *et al.*, 2019) | *F. virguliforme* | 27 |
| (Vinholes *et al.*, 2019) | *M. phaseolina* | 4 |
| (Coser *et al.*, 2017) | *M. phaseolina* | 12 |
| (Rincker *et al.*, 2016) | *C. gregata* | 7 |

### 2.2 Transcriptome data

Gene expression estimates in transcripts per million mapped reads (TPM, Kallisto estimation) were retrieved from the Soybean Expression Atlas (Machado *et al*., 2020). Additional RNA-seq samples comprising soybean tissues infected with fungal pathogens were retrieved from a recent publication from our group (Almeida-Silva & Venancio, 2021a). We filtered the SNP and transcriptome datasets to keep only fungal species that were represented by both data sources. A total of 150 RNA-seq samples from soybean tissues infected with fungal pathogens were selected (Supplementary Table S2). Finally, genes with median expression values lower than 5 were excluded to attenuate noise, resulting in an 18748 *x* 150 gene expression matrix for downstream analyses.

### 2.3 Selection of guide genes

MapMan annotations for soybean genes were retrieved from the PLAZA 3.0 Dicots database (Proost *et al*., 2015). Genes assigned to defense-related pathways (*e.g*., pathogenesis-related proteins, lignin biosynthesis, oxidative stress, and phytohormone regulation) were used as guides (Supplementary Table S3).

### 2.4 Candidate gene mining and functional analyses

Gene expression data were adjusted for confounding artifacts and quantile normalized with the R package BioNERO (Almeida-Silva & Venancio, 2021b). An unsigned coexpression network was inferred with BioNERO using Pearson’s r as correlation. All genes located in a 2 Mb sliding window relative to each SNP were selected as putative candidates, as previously proposed (Brodie *et al*., 2016). Candidate genes were prioritized using the algorithm implemented in the R package cageminer (Almeida-Silva & Venancio, 2021c), with an r_pb_ threshold of 0.2 for gene significance (gene-trait correlation). Enrichment analyses were also performed with BioNERO, using functional annotations from the PLAZA 4.0 database (Van Bel *et al*., 2018). To rank the prioritized candidates, they were given scores using the formula:

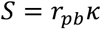

where

*r*_*pb*_ = point-biserial correlation coefficient (cageminer algorithm)

*κ* = 2 if the gene is a transcription factor

*κ* = 2 if the gene is a hub

*κ* = 3 if the gene is a hub and a transcription factor

*κ* = 1 if the gene is neither a hub nor a transcription factor

### 2.5 Selection of most resistant accessions from the USDA germplasm

The VCF file with genotypic information for all accessions in the USDA germplasm was downloaded from Soybase (Brown *et al*., 2020). Scores 0, 1, and 2 were attributed to accessions with 0, 1, and 2 beneficial SNPs (effect size >0), respectively, whereas scores 2, 1, and 0 were attributed to accessions with 0, 1, and 2 deleterious SNPs (effect size <0). The resistance potential of the best accessions was calculated as a ratio of the attributed scores to the theoretical maximum score (all beneficial SNPs and no deleterious SNPs).

## 3 Results and discussion

### 3.1 Data summary and genomic distribution of SNPs

After filtering the datasets to keep only fungal species represented by both SNP and transcriptome information, we kept five common phytopathogenic fungi: *Cadophora gregata, Fusarium graminearum, Fusarium virguliforme, Macrophomina phaseolina*, and *Phakopsora pachyrhizi* (Figure 1A). Overall, SNPs were located in gene-rich regions of the genome (Figure 1B). SNPs were unevenly distributed across chromosomes, except for *F. virguliforme* (Figure 1C). Further, we found that most SNPs were located in intergenic regions (Figure 1D). Hence, predicting SNP effect on genes would not be suitable for this trait.

**Figure 1.**
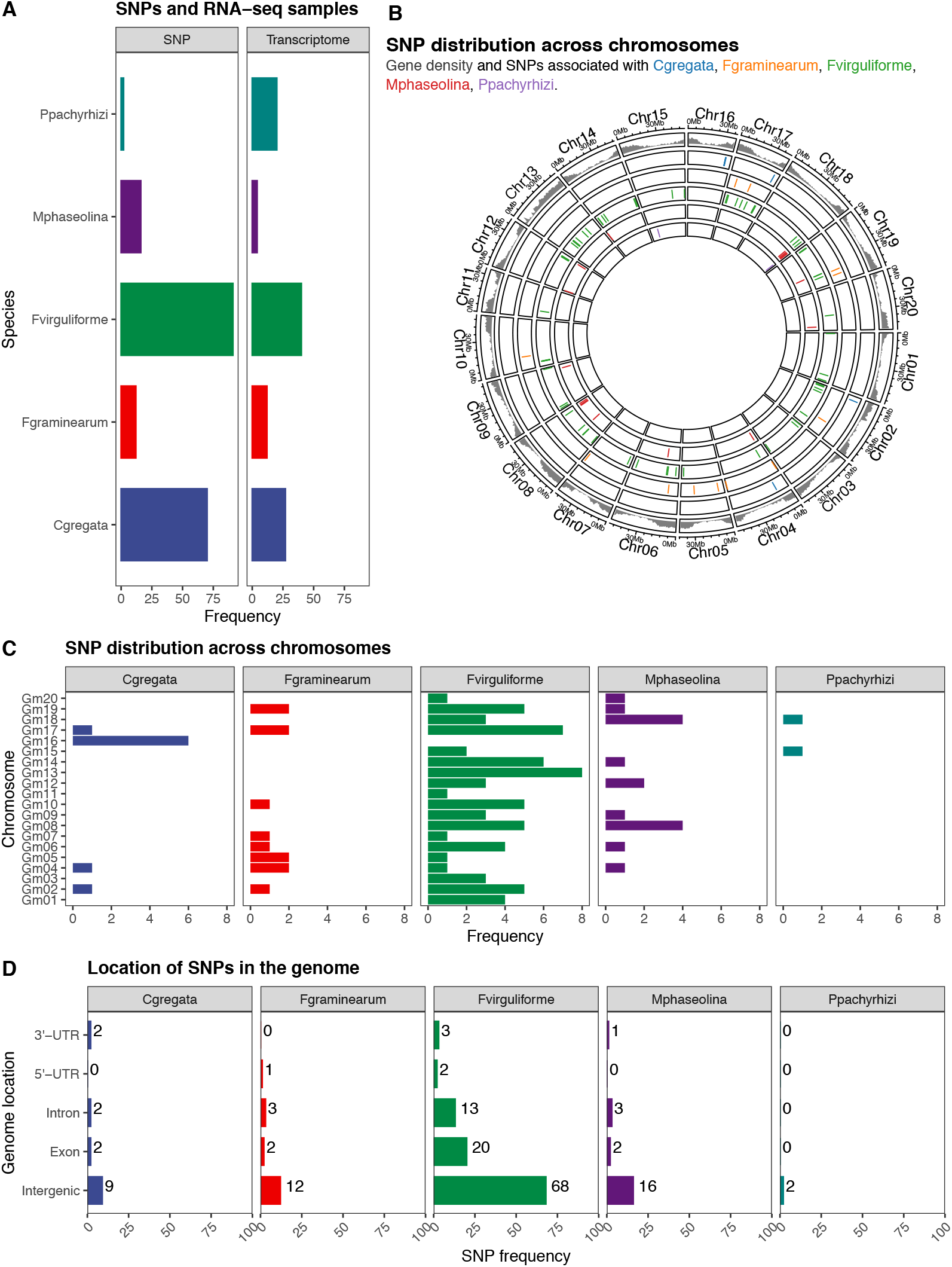
Data summary and genomic distribution of SNPs. A. Frequency of SNPs and RNA-seq samples included in this study. B. Genomic coordinates of resistance SNPs against each fungal pathogen. The outer track represents gene density, whereas inner tracks represent the SNP positions for each species. C. SNP distribution across chromosomes. Overall, there is an uneven distribution of SNPs across chromosomes. D. Genomic location of SNPs. Most SNPs are located in intergenic regions.

### 3.2 Candidate gene mining reveals a highly species-specific immune response

Using defense-related genes as guides, the cageminer algorithm identified 188, 56, 11, 8, and 3 high-confidence genes for *F. virguliforme, F. graminearum, C. gregata, M. phaseolina, and P. pachyrhizi*, respectively (Figure 2B). Only three genes were shared between species, revealing a high specificity in plant-pathogen interactions for these species. The three genes are shared by *F. virguliforme* and *F. graminearum*, suggesting that some conservation can occur at the genus level, but not at other broader taxonomic levels.

**Figure 2.**
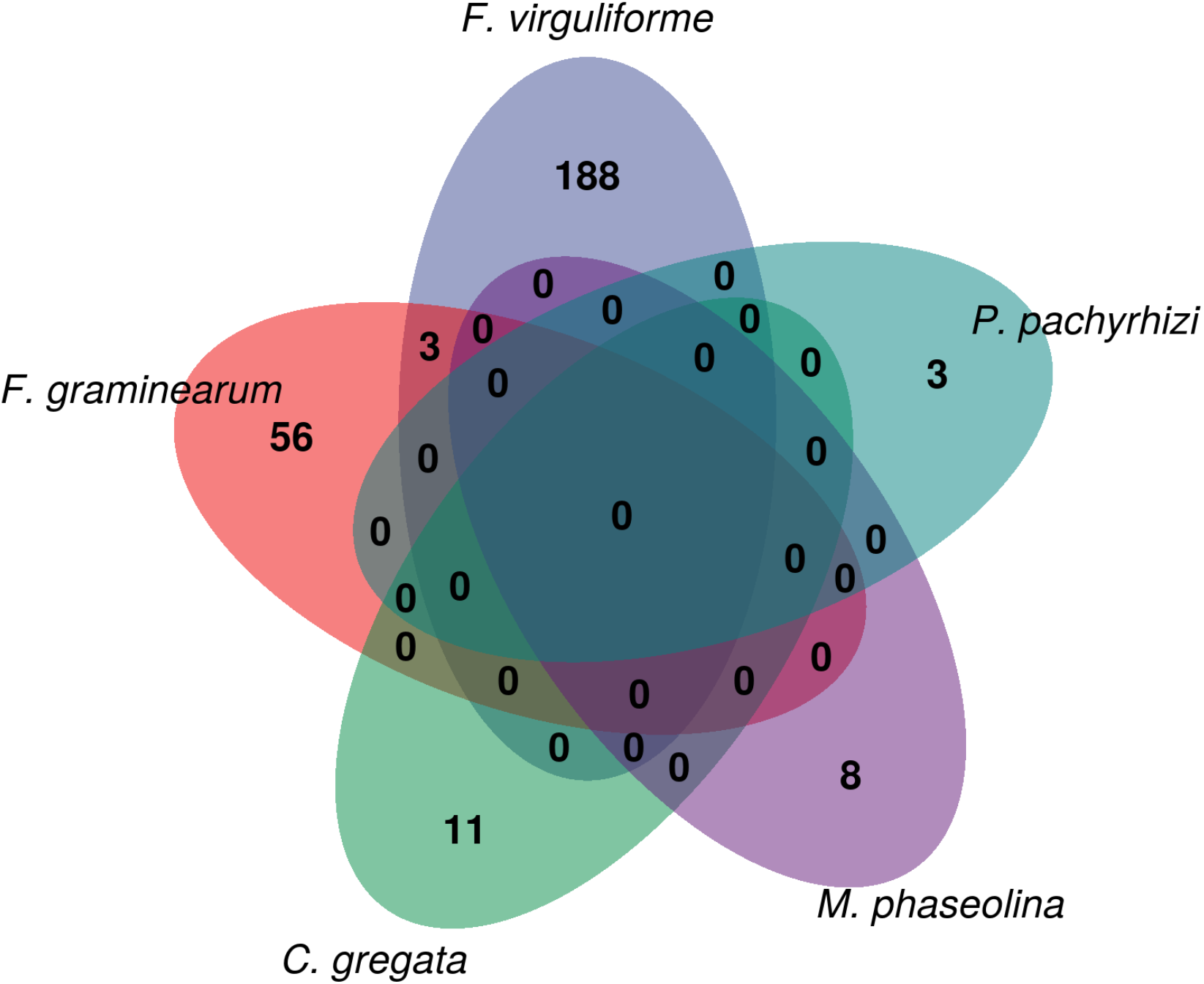
Venn diagram of prioritized candidate resistance genes against each species. The diagram demonstrates a high species-specific response to each pathogen, as genes are mostly not shared. Only three genes are shared between *F. graminearum* and *F. virguliforme*, suggesting some conservation at the genus level.

The specificity of resistance genes to particular species has been widely reported (Durrant & Dong, 2004; Kourelis & Van Der Hoorn, 2018; Ning & Wang, 2018; Li *et al*., 2020). This phenomenon imposes a challenge for biotechnological applications, as it requires pyramiding many different genes to render elite cultivars resistant to different pathogens. However, we cannot rule out that the species-specific trend we observed results from low diversity in the association panels in the GWAS we analyzed. Additionally, as SNP and transcriptome data are not available for multiple pathogen strains, we might overlook broad-spectrum resistance genes that confer resistance to multiple strains of the same species (Ning & Wang, 2018).

Further, we manually curated the high-confidence candidate resistance genes to predict the putative role of their products in plant immunity (Supplementary Table S4). Most of the prioritized candidates (28%) encode proteins involved in immune signaling, although it does not apply to all fungi species (Figure 3). Candidates also encode proteins that play a role in recognition, phytohormone metabolism, systemic acquired resistance, transport, transcriptional regulation, oxidative stress, apoptosis, physical defense, and direct function against fungi (Figure 3). Interestingly, 21 candidate genes lack functional description and, hence, we could not infer their roles in plant immunity (*n=*2, 4, 14, and 1 for *C. gregata, F. virguliforme*, and *P. pachyrhizi*, respectively). Nevertheless, as they were identified as high-confidence candidate genes, we hypothesize that they encode defense-related proteins. We also developed a scheme that was used to rank high-confidence candidate genes, which can be used to prioritize candidates for experimental validation in future studies (Table 2).

**Figure 3.**
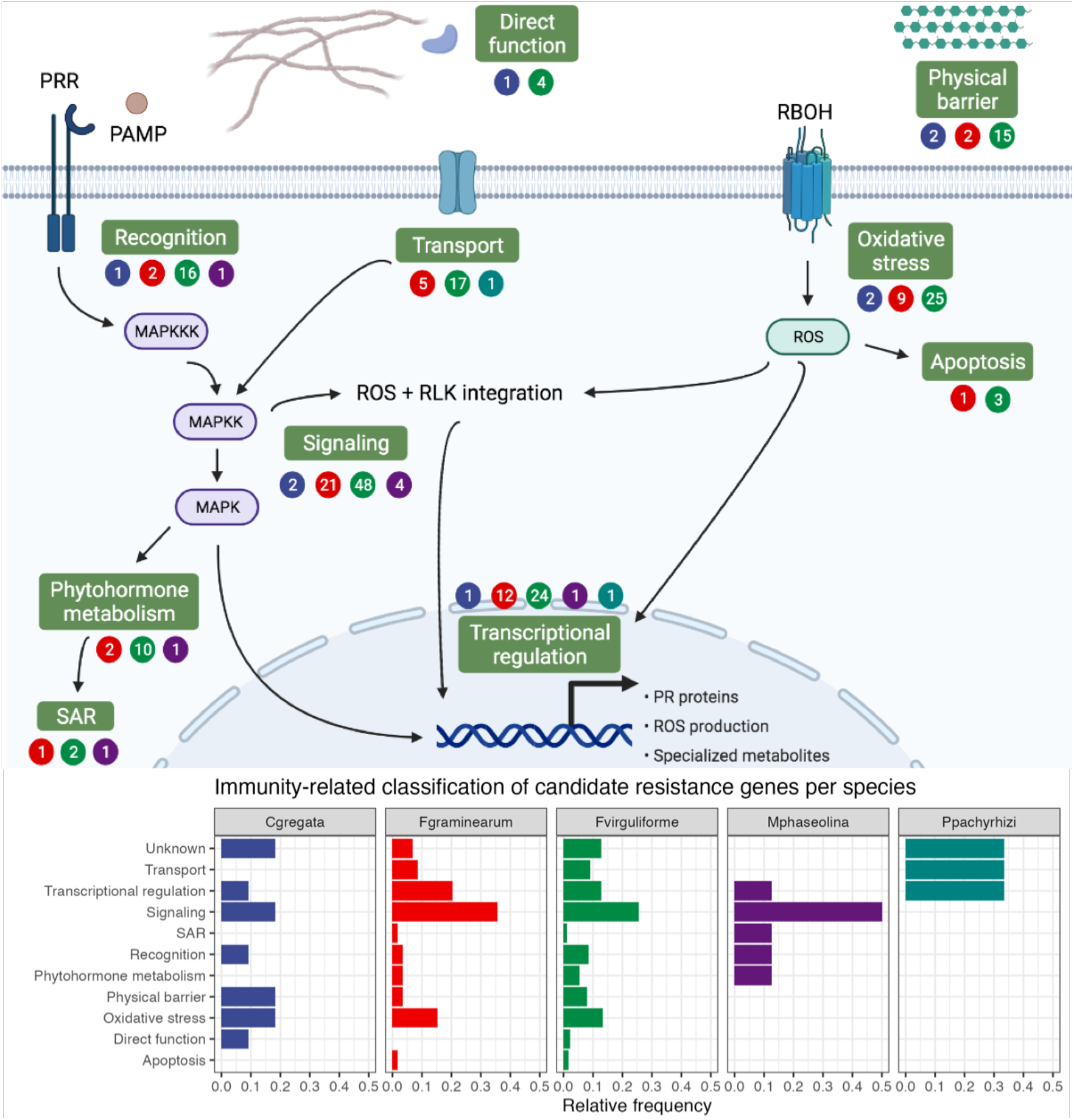
Prioritized candidate resistance genes and their putative role in plant immunity. Numbers in circles represent absolute frequencies of resistance genes against *C. gregata* (blue), *F. graminearum* (red), *F. virguliforme* (green), *M. phaseolina* (purple), and *P pachyrhizi* (turquoise). PRR, pattern recognition receptor. PAMP, pathogen-associated molecular pattern. MAPKKK, mitogen-activated protein kinase kinase kinase. MAPKK, mitogen-activated protein kinase kinase. MAPK, mitogen-activated protein kinase. SAR, systemic acquired resistance. RBOH, respiratory burst oxidase homolog. ROS, reactive oxygen species. RLK, receptor-like kinase. PR, pathogenesis-related. Figure designed with Biorender (biorender.com).

**Figure 4.**
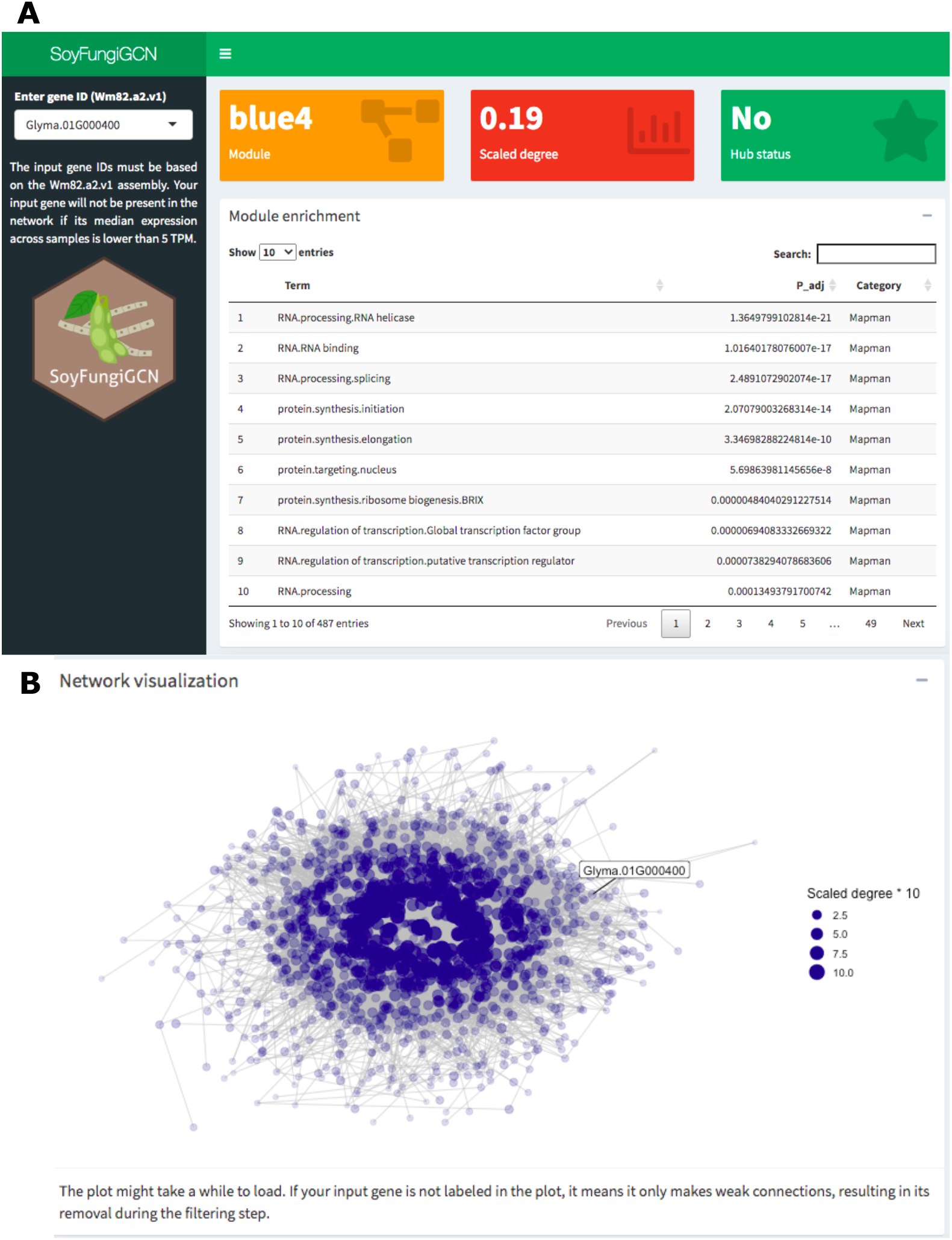
Functionalities in the SoyFungiGCN web application. A. Screenshot of the page users see when they access the application. In the sidebar, users can specify the ID of a gene of interest (Wm82.a2.v1 assembly). For each gene, users can see the gene’s module (orange box), scaled degree (red box), hub gene status (green box), and an interactive table with enrichment results for MapMan bins, Interpro domains and Gene Ontology terms associated the gene’s module. P-values from enrichment results are adjusted for multiple testing with Benjamini-Hochberg correction. B. Network visualization plot. Users can optionally visualize the input gene and its position in the module by clicking the plus (+) icon in the “Network visualization” tab below the enrichment table. As the plot can take a few seconds to render (∼2-5 seconds), it is hidden by default.

**Table 2.** Top 10 candidate resistance genes against each fungal species and their putative roles in plant immunity. The predicted function for each gene was manually curated from the description of the best ortholog in *Arabidopsis thaliana*, using functional annotations from Soybase and TAIR.

| Gene | Predicted function | Resistance to | Role |
| --- | --- | --- | --- |
| Glyma.16G170100 | Cell wall biogenesis-related extensin 3 | *C. gregata* | Physical barrier |
| Glyma.02G026700 | Transcriptional repressor SIN3 | *C. gregata* | Transcriptional regulation |
| Glyma.02G026900 | Galacturonosyltransferase | *C. gregata* | Physical barrier |
| Glyma.02G029300 | SAM domain-containing | *C. gregata* | Unknown |
| Glyma.16G155100 | Aquaporin | *C. gregata* | Oxidative stress |
| Glyma.17G217000 | Class V chitinase | *C. gregata* | Direct function |
| Glyma.17G213600 | Calcium-binding EF hand | *C. gregata* | Signaling |
| Glyma.17G231800 | Clathrin adaptor EPSIN1 | *C. gregata* | Recognition |
| Glyma.02G047000 | Thiosulfate sulfurtransferase/rhodanese | *C. gregata* | Oxidative stress |
| Glyma.16G150500 | Unknown | *C. gregata* | Unknown |
| Glyma.17G087500 | SOUL heme-binding protein | *F. graminearum* | Oxidative stress |
| Glyma.06G121300 | GRAS transcription factor | *F. graminearum* | Transcriptional regulation |
| Glyma.05G070300 | Tobamovirus multiplication 2A | *F. graminearum* | Recognition |
| Glyma.04G013500 | BURP domain-containing protein | *F. graminearum* | Physical barrier |
| Glyma.06G105000 | ERF/AP2 transcription factor | *F. graminearum* | Transcriptional regulation |
| Glyma.05G062400 | 2OG-Fe(II) oxygenase | *F. graminearum* | Oxidative stress |
| Glyma.05G063600 | ERF/AP2 transcription factor | *F. graminearum* | Transcriptional regulation |
| Glyma.05G115700 | RING domain ubiquitin E3 ligase | *F. graminearum* | Signaling |
| Glyma.17G116100 | MAPK signaling-related protein | *F. graminearum* | Signaling |
| Glyma.05G103600 | Peroxidase | *F. graminearum* | Oxidative stress |
| Glyma.13G081000 | Nodulin-like amino acid transporter | *F. virguliforme* | Transport |
| Glyma.01G225600 | Unknown | *F. virguliforme* | Unknown |
| Glyma.02G210500 | bHLH transcription factor | *F. virguliforme* | Transcriptional regulation |
| Glyma.01G162500 | BIG1 protein | *F. virguliforme* | Apoptosis |
| Glyma.17G061400 | Peroxidase | *F. virguliforme* | Oxidative stress |
| Glyma.19G010100 | HD-Zip transcription factor | *F. virguliforme* | Transcriptional regulation |
| Glyma.18G276800 | Amino acid transporter | *F. virguliforme* | Oxidative stress |
| Glyma.05G209900 | PLAC8 family protein | *F. virguliforme* | Apoptosis |
| Glyma.14G025100 | Inositol-1,4,5-trisphosphate 5-phosphatase | *F. virguliforme* | Signaling |
| Glyma.19G117800 | Unknown | *F. virguliforme* | Unknown |
| Glyma.20G203900 | Type I serine/threonine protein phosphatase | *M. phaseolina* | Signaling |
| Glyma.08G316500 | Calmodulin-dependent protein kinase | *M. phaseolina* | Signaling |
| Glyma.06G187200 | R-gene-mediated resistance, lipase | *M. phaseolina* | SAR |
| Glyma.09G218600 | Cytochrome P450, family 707, subfamily A | *M. phaseolina* | Phytohormone metabolism |
| Glyma.09G216800 | Pectin acetylesterase | *M. phaseolina* | Signaling |
| Glyma.20G216600 | Dof-type transcription factor | *M. phaseolina* | Transcriptional regulation |
| Glyma.08G332800 | Calcineurin B-like calcium sensor | *M. phaseolina* | Signaling |
| Glyma.18G301700 | Leucine-rich repeat receptor kinase (LRR-RK) | *M. phaseolina* | Recognition |
| Glyma.15G125900 | Magnesium transporter CorA-like | *P. pachyrhizi* | Transport |
| Glyma.18G286900 | Unknown | *P. pachyrhizi* | Unknown |
| Glyma.15G123900 | CBF1 interacting co-repressor CIR | *P. pachyrhizi* | Transcriptional regulation |

### 3.3 Pangenome presence/absence variation analysis demonstrates that most prioritized genes are core genes

We analyzed PAV patterns for our prioritized candidate genes in the recently published pangenome of cultivated soybeans to unveil which soybean genotypes contain prioritized candidate genes and explore gene presence/absence variation patterns across genomes (Torkamaneh *et al*., 2021). We found that most candidates are present in all 204 accessions (Supplementary Figure 1A). This trend is not surprising, as the gene content in this pangenome is highly conserved, with ∼91% of the genes being shared by >99% of the genomes. Although the variable genome is enriched in genes associated with defense, signaling, and plant development, this trend was not found in our gene set.

Further, we investigated if gene PAV patterns could be explained by the geographical origins of the accessions (Supplementary Figure 1B). Strikingly, we observed no clustering by geographical origin, suggesting that gene PAV is not affected by population structure. As this pangenome is comprised of improved soybean accessions (Torkamaneh *et al*., 2021), the lack of population structure effect can be due to breeding programs targeting optimal adaptation to different environmental conditions (*e.g*., latitude and climate), even if they are in the same country.

### 3.4 Screening of the USDA germplasm reveals a room for genetic improvement

We inspected the USDA germplasm to find the top 5 most resistant genotypes against each fungal pathogen (see Materials and Methods for details). Strikingly, the most resistant genotypes do not contain all resistance alleles, revealing that, theoretically, they could be further improved to increase resistance (Table 3). All resistance-associated SNPs against *P. pachyrhizi* are present in some accessions, but this is because only two SNPs have been reported for this species. Additionally, none of the reported SNPs for *F. graminearum* have been identified in the SoySNP50k collection. Hence, we could not predict the most resistant accessions to this fungal species in the USDA germplasm.

**Table 3.** Top 5 most resistant soybean accessions against each fungal pathogen. Overall, the best genotypes do not reach the maximum potential. An exception is observed for *P. pachyrhizi*-resistant genotypes, but this is likely due to the small number of resistance SNPs. None of the resistance SNPs for *F. graminearum* have been identified in the USDA SoySNP50k compendium and, hence, we could not predict resistance potential against this species.

| Accession | Score | Potential | Species |
| --- | --- | --- | --- |
| PI594466 | 102 | 0.73 | *C. gregata* |
| PI578477A | 100 | 0.71 | *C. gregata* |
| PI437571 | 100 | 0.71 | *C. gregata* |
| PI567520A | 100 | 0.71 | *C. gregata* |
| PI274507 | 100 | 0.71 | *C. gregata* |
| PI339871C | 82 | 0.60 | *F. virguliforme* |
| PI378694 | 80 | 0.59 | *F. virguliforme* |
| PI407145 | 80 | 0.59 | *F. virguliforme* |
| PI424107A | 80 | 0.59 | *F. virguliforme* |
| PI479753A | 80 | 0.59 | *F. virguliforme* |
| PI594760B | 24 | 0.75 | *M. phaseolina* |
| PI479752 | 24 | 0.75 | *M. phaseolina* |
| PI603706A | 24 | 0.75 | *M. phaseolina* |
| PI603531A | 24 | 0.75 | *M. phaseolina* |
| PI603412A | 24 | 0.75 | *M. phaseolina* |
| PI603547 | 4 | 1 | *P. pachyrhizi* |
| PI639559A | 4 | 1 | *P. pachyrhizi* |
| PI639559B | 4 | 1 | *P. pachyrhizi* |
| PI326582A | 4 | 1 | *P. pachyrhizi* |
| PI407057 | 4 | 1 | *P. pachyrhizi* |

Although some individual genes can confer full race-specific resistance to some pathogens, their durability in the field is often short because of pathogen evolution (Ning & Wang, 2018). Thus, pyramiding quantitative trait loci (QTL) that confer partial resistance has been proposed a strategy to confer long-term resistance (Li *et al*., 2020). To accomplish this, the most resistant genotypes identified here can be targets of allele pyramiding in breeding programs using marker-assisted selection. Alternatively, these genotypes might have their genomes edited with CRISPR/Cas systems to introduce beneficial alleles or remove deleterious alleles, ultimately boosting resistance.

### 3.5 Development of a user-friendly web application for network exploration

To facilitate network exploration and data reuse, we developed a user-friendly web application named SoyFungiGCN (https://soyfungigcn.venanciogroup.uenf.br/). Users can input a soybean gene of interest (Wm82.a2.v1 assembly) and visualize the gene’s module, scaled intramodular degree, and hub status (Figure 1A). Additionally, users can explore enriched GO terms, Mapman bins and/or Interpro domains associated with the input gene’s module (Figure 1A). Users can also visualize a network plot with the input gene and its coexpression neighbors (Figure 1B). This resource can be particularly useful for researchers studying soybean response to other fungal species, as they can check if their genes of interest are located in defense-related coexpression modules. Also, researchers studying other species can verify if the soybean ortholog of their genes of interest is located in a defense-related module. The application is also available as an R package named SoyFungiGCN (https://github.com/almeidasilvaf/SoyFungiGCN). This package lets users run the application locally as a Shiny app, ensuring the application will always be available, even in case of server downtime.

## 4 Conclusions

By integrating publicly available GWAS and RNA-seq data, we found promising candidate genes in soybean associated with resistance to five common phytopathogenic fungi, namely *C. gregata, F. graminearum, F. virguliforme, M. phaseolina*, and *P. pachyrhizi*. The prioritized candidates encode proteins that play a role immunity-related processes such as in recognition, signaling, transcriptional regulation, oxidative stress, and physical defense. We have also found the top 5 most resistant soybean accessions against each fungal species and hypothesize that they can be further genetically improved in breeding programs with marker-assisted selection or through genome editing. The coexpression network generated here was also made available in a web resource and R package to help in future studies on soybean-pathogenic fungi interactions.

## Data availability

All data and code used in this study are available in our GitHub repository (https://github.com/almeidasilvaf/SoyFungi_GWAS_GCN) to ensure full reproducibility.

## Acknowledgements

This work was supported by Fundação Carlos Chagas Filho de Amparo à Pesquisa do Estado do Rio de Janeiro (FAPERJ; grants E-26/203.309/2016 and E-26/203.014/2018), Coordenação de Aperfeiçoamento de Pessoal de Nível Superior - Brasil (CAPES; Finance Code 001), and Conselho Nacional de Desenvolvimento Científico e Tecnológico. The funding agencies had no role in the design of the study and collection, analysis, and interpretation of data and in writing.

## Author contributions

Conceived the study: FA-S and TMV. Data analysis: FA-S. Funding, project coordination and infrastructure: TMV. Manuscript writing: FA-S and TMV.

## REFERENCES

Almeida-Silva F, Venancio TM. 2021a. Pathogenesis-related protein 1 (PR-1) genes in soybean: genome-wide identification, structural analysis and expression profiling under multiple biotic and abiotic stresses. bioRxiv 1: 1–23.

Almeida-Silva F, Venancio TM. 2021b. BioNERO: an all-in-one R/Bioconductor package for comprehensive and easy biological network reconstruction. bioRxiv: 2021.04.10.439287.

Almeida-Silva F, Venancio TM. 2021c. cageminer: an R/Bioconductor package to prioritize candidate genes by integrating GWAS and gene coexpression networks. bioRxiv: 1–9.

Baker RL, Leong WF, Brock MT, Rubin MJ, Markelz RJC, Welch S, Maloof JN, Weinig C. 2019. Integrating transcriptomic network reconstruction and eQTL analyses reveals mechanistic connections between genomic architecture and Brassica rapa development. PLOS Genetics 15: e1008367.

Bandara AY, Weerasooriya DK, Bradley CA, Allen TW, Esker PD. 2020. Dissecting the economic impact of soybean diseases in the United States over two decades. PLoS ONE 15: 1–28.

Bao Y, Kurle JE, Anderson G, Young ND. 2015. Association mapping and genomic prediction for resistance to sudden death syndrome in early maturing soybean germplasm. Molecular Breeding 35: 1–14.

Baxter I. 2020. We aren’t good at picking candidate genes, and it’s slowing us down. Current Opinion in Plant Biology 54: 57–60.

Van Bel M, Diels T, Vancaester E, Kreft L, Botzki A, Van De Peer Y, Coppens F, Vandepoele K. 2018. PLAZA 4.0: An integrative resource for functional, evolutionary and comparative plant genomics. Nucleic Acids Research 46: D1190–D1196.

Brodie A, Azaria JR, Ofran Y. 2016. How far from the SNP may the causative genes be? Nucleic Acids Research 44: 6046–6054.

Brown A V, Conners SI, Huang W, Wilkey AP, Grant D, Weeks NT, Cannon SB, Graham MA, Nelson RT. 2020. A new decade and new data at SoyBase, the USDA-ARS soybean genetics and genomics database. Nucleic Acids Research 13: 1–6.

Chang HX, Lipka AE, Domier LL, Hartman GL. 2016. Characterization of disease resistance loci in the USDA soybean germplasm collection using genome-wide association studies. Phytopathology 106: 1139–1151.

Coser SM, Reddy RVC, Zhang J, Mueller DS, Mengistu A, Wise KA, Allen TW, Singh A, Singh AK. 2017. Genetic architecture of charcoal rot (Macrophomina phaseolina) resistance in soybean revealed using a diverse panel. Frontiers in Plant Science 8: 1–12.

Deshmukh R, Sonah H, Patil G, Chen W, Prince S, Mutava R, Vuong T, Valliyodan B, Nguyen HT. 2014. Integrating omic approaches for abiotic stress tolerance in soybean. Frontiers in Plant Science 5: 1–12.

Durrant WE, Dong X. 2004. Systemic acquired resistance. Annual Review of Phytopathology 42: 185–209.

Iquira E, Humira S, François B. 2015. Association mapping of QTLs for sclerotinia stem rot resistance in a collection of soybean plant introductions using a genotyping by sequencing (GBS) approach. BMC Plant Biology 15: 1–12.

Kandel R, Chen CY, Grau CR, Dorrance AE, Liu JQ, Wang Y, Wang D. 2018. Soybean resistance to white mold: Evaluation of soybean germplasm under different conditions and validation of QTL. Frontiers in Plant Science 9: 1–12.

Kourelis J, Van Der Hoorn RAL. 2018. Defended to the Nines: 25 years of Resistance Gene Cloning Identifies Nine Mechanisms for R Protein Function. Plant Cell.

Li W, Deng Y, Ning Y, He Z, Wang GL. 2020. Exploiting Broad-Spectrum Disease Resistance in Crops: From Molecular Dissection to Breeding. Annual Review of Plant Biology 71: 575–603.

Machado FB, Moharana KC, Almeida - Silva F, Gazara RK, Pedrosa - Silva F, Coelho FS, Grativol C, Venancio TM. 2020. Systematic analysis of 1,298 RNA - Seq samples and construction of a comprehensive soybean (Glycine max) expression atlas. The Plant Journal: 0–2.

Michno JM, Liu J, Jeffers JR, Stupar RM, Myers CL. 2020. Identification of nodulation-related genes in Medicago truncatula using genome-wide association studies and co-expression networks. Plant Direct 4: 1–10.

Ning Y, Wang GL. 2018. Breeding plant broad-spectrum resistance without yield penalties. Proceedings of the National Academy of Sciences of the United States of America 115: 2859–2861.

Pandey AK, Yang C, Zhang C, Graham MA, Horstman HD, Lee Y, Zabotina OA, Hill JH, Pedley KF, Whitham SA. 2011. Functional analysis of the asian soybean rust resistance pathway mediated by Rpp2. Molecular Plant-Microbe Interactions 24: 194–206.

Proost S, Van Bel M, Vaneechoutte D, Van de Peer Y, Inzé D, Mueller-Roeber B, Vandepoele K. 2015. PLAZA 3.0: an access point for plant comparative genomics. Nucleic Acids Research 43: D974–D981.

Rincker K, Lipka AE, Diers BW. 2016. Genome-Wide Association Study of Brown Stem Rot Resistance in Soybean across Multiple Populations. The Plant Genome 9: plantgenome2015.08.0064.

Schaefer RJ, Michno J-M, Jeffers J, Hoekenga O, Dilkes B, Baxter I, Myers CL. 2018. Integrating Coexpression Networks with GWAS to Prioritize Causal Genes in Maize. The Plant Cell 30: 2922–2942.

Schwartz TS. 2020. The promises and the challenges of integrating multi-omics and systems biology in comparative stress biology. Integrative and Comparative Biology 53: 1689–1699.

Sun M, Jing Y, Zhao X, Teng W, Qiu L, Zheng H, Li W, Han Y. 2020. Genome-wide association study of partial resistance to sclerotinia stem rot of cultivated soybean based on the detached leaf method. PLoS ONE 15: 1–15.

Swaminathan S, Das A, Assefa T, Knight JM, Da Silva AF, Carvalho JPS, Hartman GL, Huang X, Leandro LF, Cianzio SR, et al. 2019. Genome wide association study identifies novel single nucleotide polymorphic loci and candidate genes involved in soybean sudden death syndrome resistance. PLoS ONE 14: 1–21.

Torkamaneh D, Lemay M-A, Belzile F. 2021. The Pan-genome of the Cultivated Soybean (PanSoy) Reveals an Extraordinarily Conserved Gene Content. Plant Biotechnology Journal n/a.

Vinholes P, Rosado R, Roberts P, Borém A, Schuster I. 2019. Single nucleotide polymorphism-based haplotypes associated with charcoal rot resistance in Brazilian soybean germplasm. Agronomy Journal 111: 182–192.

Wen Z, Tan R, Zhang S, Collins PJ, Yuan J, Du W, Gu C, Ou S, Song Q, An YQC, et al. 2018. Integrating GWAS and gene expression data for functional characterization of resistance to white mould in soya bean. Plant Biotechnology Journal 16: 1825–1835.

Zhang J, Singh A, Mueller DS, Singh AK. 2015. Genome-wide association and epistasis studies unravel the genetic architecture of sudden death syndrome resistance in soybean. Plant Journal 84: 1124–1136.

Zhang C, Zhao X, Qu Y, Teng W, Qiu L, Zheng H, Wang Z, Han Y, Li W. 2019. Loci and candidate genes in soybean that confer resistance to Fusarium graminearum. Theoretical and Applied Genetics 132: 431–441.

