## Supplementary Figures for "Integration of genome-wide association studies and gene coexpression networks unveils promising soybean resistance genes against five common fungal pathogens"

[1]Laboratório de Química e Função de Proteínas e Peptídeos, Centro de Biociências e Biotecnologia, Universidade Estadual do Norte Fluminense Darcy Ribeiro, Campos dos Goytacazes, RJ, Brazil.

*TMV: Laboratório de Química e Função de Proteínas e Peptídeos, Centro de Biociências e Biotecnologia, Universidade Estadual do Norte Fluminense Darcy Ribeiro. Av. Alberto Lamego 2000, P5, sala 217, Campos dos Goytacazes, RJ, Brazil.

*FA-S: Laboratório de Química e Função de Proteínas e Peptídeos, Centro de Biociências e Biotecnologia, Universidade Estadual do Norte Fluminense Darcy Ribeiro. Av. Alberto Lamego 2000, P5, sala 217, Campos dos Goytacazes, RJ, Brazil.

### Supplementary Figures

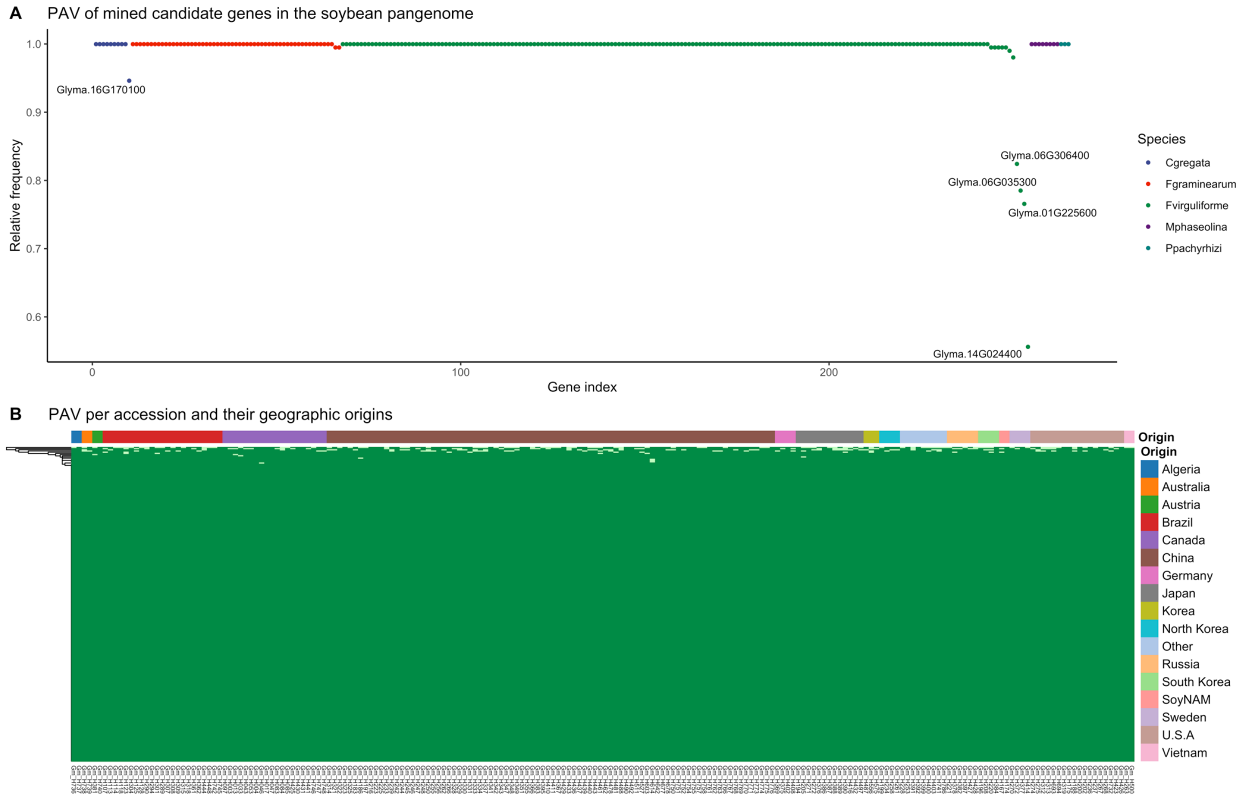


**Figure 1.** Presence/absence variation (PAV) of prioritized candidate genes in the soybean pangenome. A. Relative frequency of accessions containing each candidate gene. Most candidates are present in all accessions. Candidate genes with lower frequency in the pangenome are labeled. B. PAV per accessions and their geographic distribution. The patterns of gene PAV cannot be explained by the geographic origins of the accessions.
